# Cell-Free Protein Crystallization Enables Rapid Structure Determination of Disaccharides and Trisaccharides Using Galectin-10 Crystals

**DOI:** 10.1101/2025.07.09.663810

**Authors:** Mariko Kojima, Xinchen Yao, Satoshi Abe, Tadaomi Furuta, Kunio Hirata, Ririko Kobayashi, Taiga Suzuki, Takafumi Ueno

**Author notes:** **Corresponding Author** Takafumi Ueno. Institute of Multidisciplinary Research for Advanced Materials, Tohoku University, Katahira 2-1-1, Aoba-ku, Sendai, Miyagi, 980-8577, Japan (Mariko Kojima).

## Abstract

It is critical to understand the conformational selection and dynamics of flexible saccharides via protein-ligand interactions in efforts to elucidate their biofunctional roles. Protein crystals can serve as scaffolds to immobilize small molecules, enabling structural and dynamic analysis of saccharides that are difficult to study by conventional approaches. However, constructing versatile scaffold crystals for high-throughput structural analysis remains challenging because this work involves laborious protein production and crystallization workflows. Here, we report rapid crystallization and structural analysis of saccharide-bound scaffolds by applying cell-free protein crystallization (CFPC) to galectin-10 (Gal-10), a lectin known to crystallize spontaneously *in vivo*. Using CFPC-generated Gal-10 crystals, we obtained the first atomic-resolution structures of melezitose, one of the trisaccharide, bound to the protein scaffold, revealing binding modes inaccessible by conventional approaches. Normalized *B*-factor analysis combined with molecular dynamics simulations reveals how the binding-site architecture modulates saccharide flexibility and immobilization. This platform can be extended to other flexible ligands and fragment-based screening.

## Introduction

The conformational selection of biomolecules such as saccharides, peptides, and nucleotides, in response to the external environment is an efficient trigger for regulating biological functions.^1, 2^ Saccharides control various *in vivo* processes such as cell signaling and glycosylation via intermolecular interactions with proteins.^1–5^ Trisaccharides and disaccharides each play important roles in molecular recognition of glycoproteins due to the flexibility of saccharides which is attributed to the rotatable glycosidic linkage between monosaccharides.^6, 7^ While NMR spectroscopy and computational modeling have provided valuable insights into saccharide structures and interactions, these methods are incapable of fully capturing the dynamic and flexible nature of saccharides.^1, 3, 4, 8–10^ To understand the relationship between conformational selection and molecular function, it is crucial to develop advanced techniques that provide atomic-resolution structures of saccharides in tunable protein environments.

Scaffold protein crystals provide a promising approach to resolving the structures of flexible molecules that undergo conformational selection. High-resolution protein crystal serves as the effective scaffold to trap target molecules in a homogenous conformation by stabilizing them via interaction with surrounding amino acids.^11–15^ Such scaffold-assisted structural analysis allows for a comprehensive analysis of conformational changes of targets in response to the external environment by mutating residues of the crystal to identify key structural determinants governing the conformation.^15^ However, constructing versatile scaffold crystals for high-throughput structural analysis remains challenging because this work involves laborious protein production and crystallization workflows. To address this problem, we developed the cell-free protein crystallization (CFPC) method, which utilizes in-cell crystallizing proteins that form high-resolution crystals within living cells.^16^ This method completes the protein synthesis and crystallization within 24 hours without protein purification. In our previous studies, we successfully demonstrated the high-throughput structure determination of intrinsically disordered proteins (IDPs), which are generally difficult to crystallize.^17^ By applying this approach to saccharides, we aim to elucidate how these molecules interact with proteins under varying molecular environments.

Galectin-10 (Gal-10), a sugar-binding protein, is a promising candidate for use as a scaffold protein crystal to capture saccharide substrates.^18^ Gal-10 is a member of the galectin family that binds to β-galactosides and is known to form crystals spontaneously *in vivo,* which are called as Charcot–Leyden crystals.^19–21^ Gal-10 has been reported to bind to several monosaccharides and organic compounds, and its structure is well characterized.^22–27^ Nevertheless, Gal-10 has not been widely used as a scaffold crystal because it requires crystallization *in vivo* or recrystallization *in vitro*, both of which are time-consuming and difficult to manage. To overcome these limitations, we employed our novel method, CFPC, that allows for rapid crystallization and structural analysis without extensive condition screening.^16^

Here, we synthesized a high-resolution crystal of Gal-10 using CFPC and investigated the use of Gal-10 scaffold crystals for disaccharide and trisaccharide structure determination. Trisaccharides have attracted attention as prebiotics because they are more resistant to hydrolysis than disaccharides, and remain in the body for longer periods.^28–31^ Prebiotics stimulate the growth of gut microbes through interactions with proteins such as transporters and metabolic enzymes.^32^ Despite the high potential of trisaccharides, structural analysis of their conformational changes upon protein binding remains challenging due to their high flexibility and lower binding specificity. After immobilizing trisaccharides in Gal-10 crystals by soaking, we identified previously unreported trisaccharide structures. Next, we synthesized the mutant Gal-10 to determine the conformation of trisaccharide which differs from that in the wild-type Gal-10, allowing us to elucidate the key interactions that stabilize trisaccharides. These findings provide new insights into the binding mechanism of trisaccharides to proteins and demonstrates the broad tolerance of saccharide-binding sites in various proteins. Furthermore, the use of CFPC with Gal-10 holds great potential for protein crystal engineering, enabling the design of scaffold protein libraries which are expected to facilitate the rapid structural analysis of small molecules, including those with unknown structures.

## Results

### Crystallization of Gal-10 using CFPC

The wild-type Gal-10 was crystallized using the Wheat Germ Protein Synthesis kit (WEPRO®7240 Expression Kit), as reported previously (See the Experimental Procedures in SI).^16^ The translation reaction was carried out using the dialysis method.^33^ Gal-10 was expressed in 80 µL of the reaction mixture for 3 days. After centrifuging the translation mixture, we obtained spindle-shaped crystals with a diameter of ∼200 µm (Figure 1a). These crystals are characterized by SDS-PAGE and MALDI-TOF and designated **Gal-10_cf_** (See SI and Figure S1).

**Figure 1.**
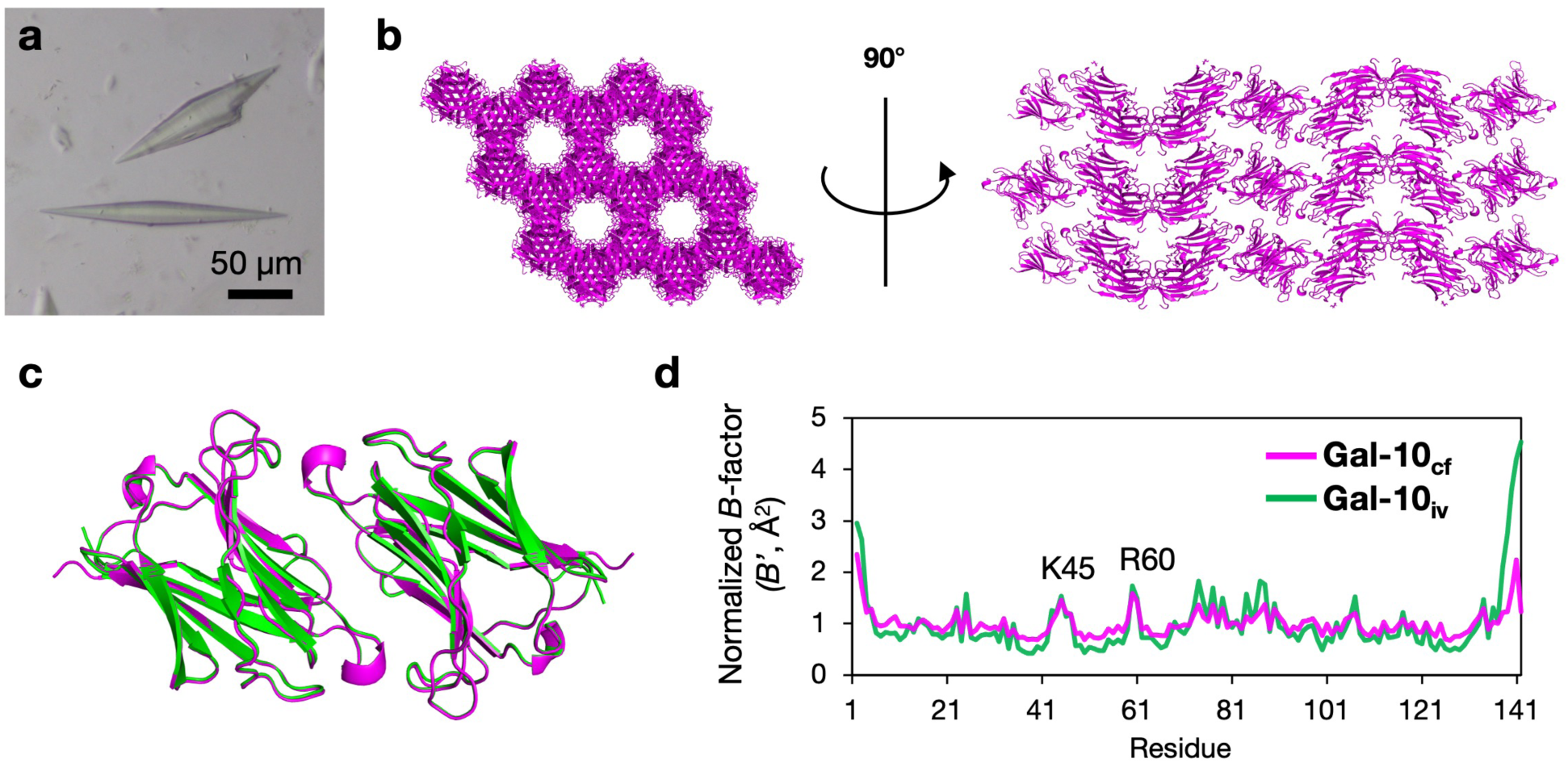
Structure analysis of **Gal-10_cf_**. (a) Microscopic image of **Gal-10**_cf_. (b) Lattice structure of **Gal-10_cf_**. (c) Dimeric structures of **Gal-10_cf_** (magenta) and Gal-10_iv_ (green). (d) Normalized per-residue *B*-factors (*B’*) of **Gal-10_cf_** and Gal-10_iv_. All proteins are shown as ribbon models.

### Structural determination of Gal-10_cf_

The crystal structures of **Gal-10_cf_** were determined at 1.60 Å resolution using BL32XU at SPring-8 (Figure 1b and Table S1). This structure has the same space group (*P*6_5_22) and the highly homologous lattice constants as the Gal-10 crystals synthesized *in vitro* (PDB ID: 1LCL, designated Gal-10_iv_) (Figure 1b). The all-atom root-mean-square deviation (RMSD) value of **Gal-10_cf_** from Gal-10_iv_ is 0.240 Å. The main chain structure of **Gal-10_cf_** is identical to that of Gal-10_iv_ (Figure 1c). The normalized per-residue *B*-factors (*B*′), index of the relative flexibility of residues within a protein structure, of **Gal-10_cf_** are similar to that of Gal-10_iv_, with local peaks at residues K45 and R60, which are likely due to the flexibility of their side chains (Figure 1d, See SI).

### Structure analysis of trisaccharide and disaccharide bound Gal-10_cf_

We attempted to determine the structures of two trisaccharides (melezitose and raffinose) and three disaccharides (sucrose, trehalose, and maltose) bound to **Gal-10_cf_**, as there were no reports on the binding structures of these saccharides with Gal-10. In addition, as a control structure, we also attempted to determine the structure of lactose in Gal-10 for a comparison with the reported structure of lactose bound to the E33A mutant of Gal-10 (E33A-Gal-10_iv_).^26^ To immobilize these saccharides into **Gal-10_cf_**, the crystals were each soaked in aqueous solutions of melezitose (1 M), raffinose (the saturated concentration, ca. 0.375 M), sucrose (1 M), trehalose (1 M), maltose (1 M), and lactose (the saturated concentration, ca. 0.6 M) at 20℃ for 24 h (See the Experimental Procedures in SI). These are designated **Mel/Gal-10_cf_**, **Raf/Gal-10_cf_**, **Suc/Gal-10_cf_**, **Tre/Gal-10_cf_**, **Mal/Gal-10_cf_**, and **Lac/Gal-10_cf_**, respectively. All crystal structures were determined at 1.57–2.31 Å resolution using BL32XU at SPring-8 (Table S1). The binding structures of the saccharides were determined from the Fourier difference maps (Figure 2 and S2).

**Figure 2.**
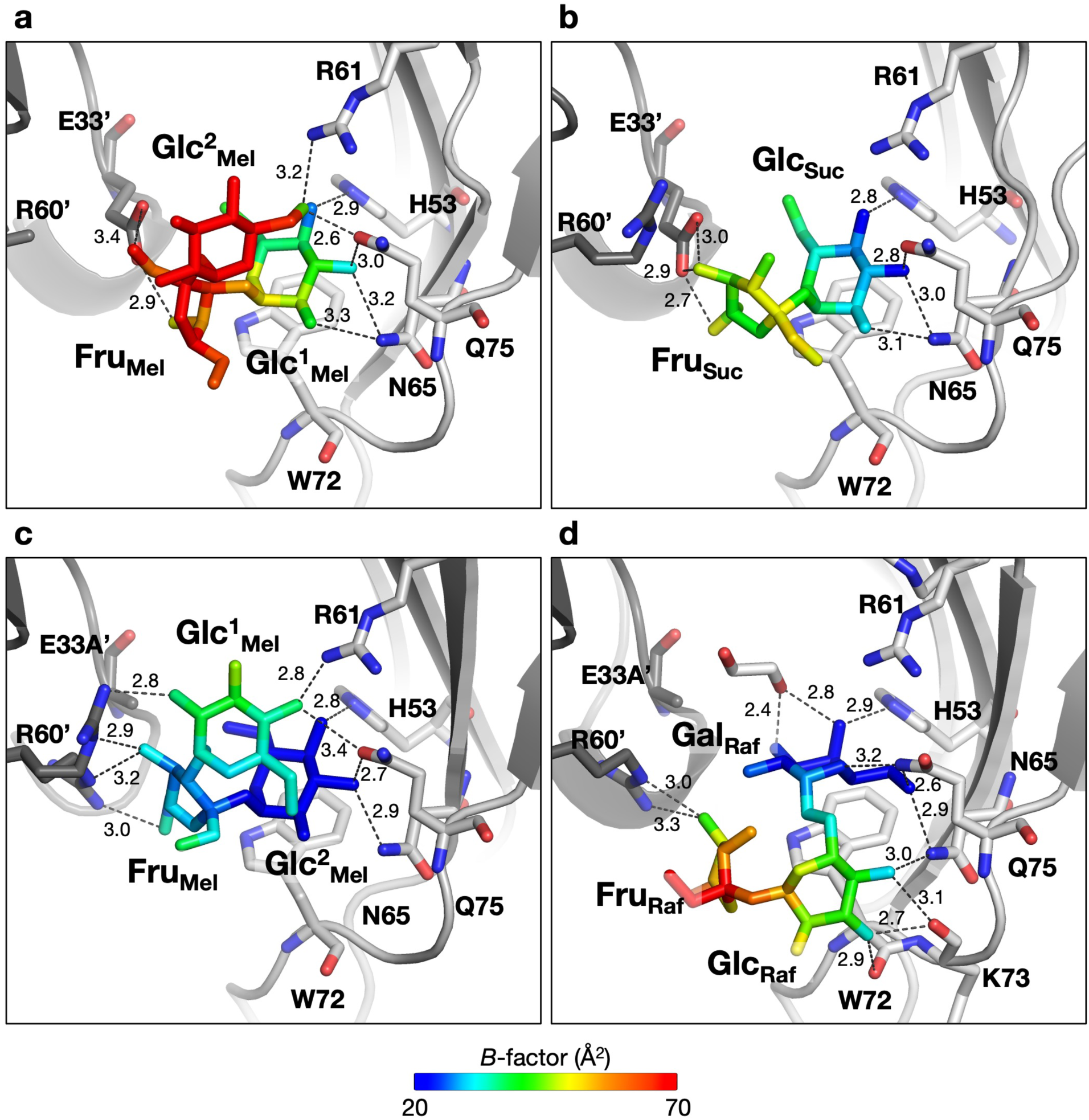
Crystal structures and the noncovalent interaction networks of (a) **Mel/Gal-10_cf_**, (b) **Suc/Gal-10_cf_**, (c) **Mel/E33A-Gal-10_cf_**, and (d) **Raf/E33A-Gal-10_cf_**. Saccharides are shown as stick models colored by the *B*-factor with values ranging from 20 Å^2^ to 70 Å^2^. Proteins are shown as ribbon models. The structure of proteins is shown as gray-colored ribbon and stick models. The cut-off distance of OH---O and NH---O hydrogen bonds is 3.5 Å.^34^

The structure we obtained for **Mel/Gal-10_cf_** is the first known example of determination of a crystal structure of a protein-melezitose complex. The structure of one of the glucose units of melezitose (Glc^1^_Mel_) was found to have a *B*-factor of 39.66 Å^2^ (Figure 2a and S2a). The *B*-factors of the other glucose unit (Glc^2^_Mel_) and the fructose unit (Fru_Mel_) are 77.03 Å^2^ and 63.30 Å^2^, respectively, suggesting that these saccharide units are less fixed than Glc^1^_Mel_. Glc^1^_Mel_ is stabilized by aromatic interactions with W72 and hydrogen bonds at O2/Glc^1^_Mel_–Nδ2/N65 (3.3 Å), O3/Glc^1^_Mel_–Nδ2/N65 (3.2 Å), O3/Glc^1^_Mel_–Oε1/Q75 (3.0 Å), and O4/Glc^1^_Mel_–Nε2/H53 (2.9 Å) (Figure 2a). These bond lengths are similar in the complex of glucose bound to Gal-10 (Glc/Gal-10_iv_) (Figure S3).^27^ In **Raf/Gal-10_cf_**, raffinose could not be determined due to a lack of the corresponding electron density.

All disaccharides were observed at the saccharide-binding site of **Gal-10_cf_**. In **Suc/Gal-10_cf_**, the whole structure of sucrose was determined (Figure 2b and S2b). The average *B*-factor of sucrose has the smallest value (32.97 Å^2^) among the four disaccharides (Table S1). The *B*-factors of glucose (Glc_Suc_) and fructose (Fru_Suc_), which are the monosaccharide units of sucrose, are 28.30 Å^2^ and 38.29 Å^2^, respectively (Table S1), indicating that Glc_Suc_ is more conformationally restrained in the crystal structure than Fru_Suc_. This Glc_Suc_ unit is stabilized by hydrogen bonds at Gal-10, O2/Glc_Suc_^1^–Nδ2/N65 (3.1 Å), O3/Glc_Suc_^1^–Nδ2/N65 (3.0 Å), O3/Glc^1^_Suc_–Oε1/Q75 (2.8 Å), and O4/Glc^1^_Suc_–Nε2/H53 (2.8 Å) along with aromatic interactions with W72 (Figure 2b). The conformation of Glc_Suc_ is identical to that of Glc^1^_Mel_. Fru_Suc_ is stabilized by hydrogen bonds at O4/Fru_Suc_–Oε1/E33’ (3.0 Å), O4/Fru_Suc_–Oε2/E33’ (2.9 Å), and O6/Fru_Suc_–Oε2/E33’ (2.7 Å) (Figure 2b). This is the first known example of a crystal structure of fructose unit bound to Gal-10.

The structures of other disaccharides were determined with average *B*-factors ranging from 39.34 Å^2^ to 52.40 Å^2^, suggesting that these disaccharides exhibit lower conformational stability than sucrose in the crystal (Figure S2c-e and Table S1). In **Tre/Gal-10_cf_** and **Mal/Gal-10_cf_**, Glc^1^_Tre_ and Glc^1^_Mal_ are fixed in a different conformations from that of Glc_Suc_ (Figure 2b, S4a, and S4b). The six-membered ring of Glc^1^_Tre_ is rotated 60° relative to that of Glc_Suc_ although both Glc^1^_Tre_ and Glc_Suc_ contain the same glucose unit with a glycosidic linkage at O1. Glc_Tre_ is stabilized by the hydrogen bonds of O1/Glc^1^_Tre_–Nδ2/N65 (3.3 Å), O2/Glc^1^_Tre_–Nδ2/N65 (3.1 Å), O2/Glc^1^_Tre_– Oε1/Q75 (2.9 Å), and O3/Glc^1^_Tre_–Νε2/H53 (2.7 Å) (Figure S4a). While the rotation of Glc^1^_Tre_ induces formation of hydrogen bonds at O6/Glc^1^_Tre_–Nη1/R60’ (3.0 Å) and O2/Glc^2^_Tre_–Oε1/Q75 (3.5 Å), that are not formed in **Suc/Gal-10_cf_**, these hydrogen bonds are weak compared to the other interactions of trehalose as indicated by the high *B*-factor (Figure 2b and S4a). In **Mal/Gal-10_cf_**, Glc^1^_Mal_ is fixed in the binding pocket via hydrogen bonds with H53, Q75, N65, E33’, and R60’ (Figure S4b). This result suggests that glucose units with glycosidic bonds at O4 preferentially bind to the binding pocket rather than those with glycosidic bonds at O1. In **Lac/Gal-10_cf_,** the structure of lactose was successfully determined, in contrast to prior experimental results indicating that the structure of lactose could only be determined in E33A-Gal-10 (Figure S4c). The orientation of lactose of **Lac/Gal-10_cf_** differs from that of the complex E33A-Gal-10_iv_ (Lac/E33A-Gal-10_iv_) (Figure S5).^26^ Deletion of the E33’ side chain induces a conformational change of R60’, which alters the shape of the saccharide-binding site. The orientation of lactose is changed to enable it to enter the cavity and the hydrogen bond pairs are optimized. This result suggests that R60’ contributes to fixation of the orientation of lactose in the crystal by modifying the shape of the saccharide-binding site rather than by formation of noncovalent bonds as indicated in the previous report.^26^

### Immobilization of trisaccharide in E33A-Gal-10_cf_

Next, we attempted to determine the structure of trisaccharide in E33A-Gal-10. E33A-Gal-10 was reported to capture disaccharide and stabilize its structure in the crystal in an arrangement which is appropriate for structure determination.^26^ **E33A-Gal-10_cf_** was synthesized using the same methods as **Gal-10_cf_** and soaked in melezitose (1 M) and raffinose (0.375 M), to produce complexes which are designated **Mel/E33A-Gal-10_cf_** and **Raf/E33A-Gal-10_cf_**, respectively (See the Experimental Procedures in SI). The crystal structures of **Mel/E33A-Gal-10_cf_** and **Raf/E33A-Gal-10_cf_** were determined at 1.62 Å and 1.57 Å resolution, respectively (Table S1).

The structures of melezitose and raffinose in **E33A-Gal-10_cf_** were successfully determined (Figure 2c,d and S2f,g). The conformation of melezitose in **E33A-Gal-10_cf_** is inverted relative to the conformation of **Mel/Gal-10_cf_** (Figure 3a). The *B*-factors of Glc^1^_Mel_, Fru_Mel_, and Glc^2^_Mel_ are 35.63 Å^2^, 30.27 Å^2^, and 16.20 Å^2^, respectively. These *B-*factors are lower than those of **Mel/Gal-10_cf_** (39.66 Å^2^, 63.30 Å^2^, and 77.03 Å^2^, respectively). The structures are fixed at Glc^2^_Mel_ and Fru_Mel_ by the hydrogen bonds with R60’, whereas the side chain becomes disordered in **Mel/Gal-10_cf_** due to steric hindrance with E33’ (Figure 2a,c). Glc^1^_Mel_ is fixed at the O2 and O4 atoms by the hydrogen bonds of O2/Glc^1^_Mel_–Nη2/R60’ (2.8 Å), O4/Glc^1^_Mel_–Nη2/R61 (2.8 Å), and O4/Glc^1^_Mel_–Oε1/Q75 (3.4 Å). Fru_Mel_ is fixed by the hydrogen bonds of O6/Fru_Mel_–Nη2/R60’ (3.0 Å) and O4/Fru_Mel_– Nε/R60’ (3.2 Å or 2.9 Å) (Figure 2c). These interactions were formed instead of the hydrogen bond interaction networks, including E33’ observed in **Mel/Gal-10_cf_**. The other interactions at Glc^2^_Mel_ are identical to those of Glc^1^_Mel_ in **Mel/Gal-10_cf_**. The structure of raffinose was determined in **Raf/E33A-Gal-10_cf_** (Figure 2d). The conformation of Gal_Raf_, the terminal monosaccharide unit fixed by W72, is similar to the conformation of Gal_Lac_ in Lac/E33A-Gal-10_iv_ (Figure 3b). The *B*-factor of Gal_Raf_ is 21.65 Å^2^. Gal_Raf_ is fixed tightly by the hydrogen bonds at O4/Gal_Raf_–Nε2/H53 (2.9 Å), O6/Gal_Raf_–Nδ2/N65 (2.9 Å), and O6/Gal_Raf_–Nε2/Q75 (2.6 Å) (Figure 2d). Gal_Raf_ has the same hydrogen bond network as Gal_Lac_ (Figure S6). Fru_Raf_, another terminal unit of raffinose, is fixed by the hydrogen bonds of O4/Fru_Raf_–Nη2/R60’ (3.3 Å) and O4/Fru_Raf_–Nε/R60’ (3.0 Å). The conformation of the side chain of R60’ is changed from the conformation in **Raf/Gal-10_cf_** to expand the space for accommodation of raffinose and to optimize the hydrogen bond network (Figure S7). These results indicate that the mutation of E33A induces hydrogen bond formation between R60’ and the terminal monosaccharide unit of a trisaccharide.

**Figure 3.**
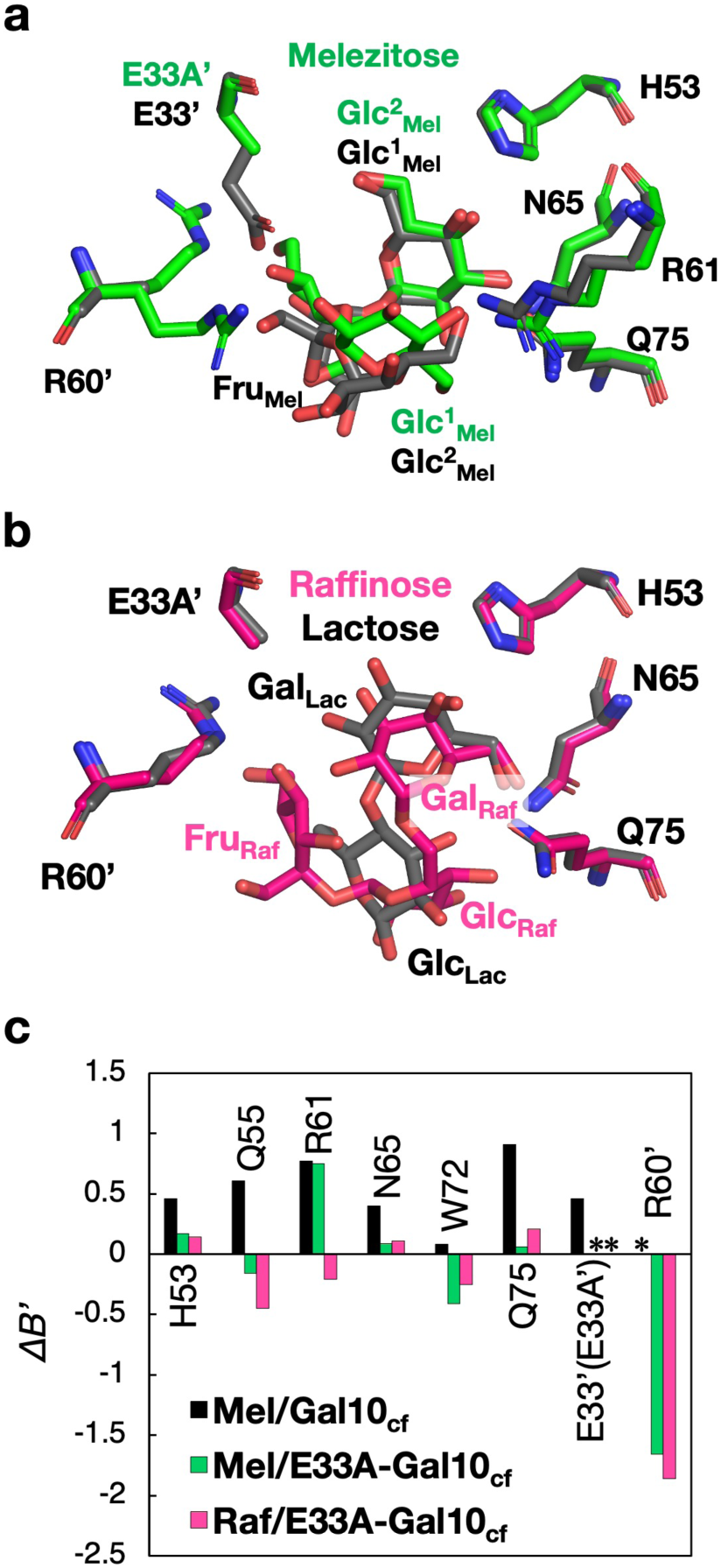
The overlayed structure of saccharide-binding sites of (a) **Mel/E33A-Gal-10_cf_** (green) and **Mel/Gal-10_cf_** (gray) and (b) **Raf/E33A-Gal-10_cf_** (pink) and **Lac/E33A-Gal-10_iv_** (gray). (c) Analysis of the difference of the normalized *B*-factor, *ΔΒ’* = *B’* – *B’_apo_*, of **Mel/ Gal-10_cf_**, **Mel/E33A-Gal-10_cf_**, and **Raf/E33A-Gal-10_cf_**.^35^ The structure of all amino acids and saccharides is shown as stick models. *No data are available due to the mutation or lack of a model of the side chain.

To compare the changes in the flexibility of residues which occur at the binding site during the transition from the unbound state to the trisaccharide-bound state, we calculated the difference in normalized *B*-factor (*11B′*, See the Experimental Procedures in SI) at the eight residues that interact with trisaccharide by side chain from ligand-free **Gal-10_cf_** to **Mel/Gal-10_cf_**, **Mel/E33A-Gal-10_cf,_** and **Raf/E33A-Gal-10_cf_** (Figure 3c). A positive value of *11B′* means that the flexibility increases after binding, and a negative value means that the flexibility decreases after binding.^35^ Among the eight residues that provide side-chain interactions with the trisaccharide, H53, Q55, R61, N65, W72, and Q75 exhibit the positive *ΔB’* values in **Mel/Gal-10_cf_**. This indicates that the flexibility of the binding site in **Mel/ Gal-10_cf_** increases after saccharide binding. While H53, R61, N65, and Q75 also exhibit positive *ΔB’* values in **Mel/E33A-Gal-10_cf_**, the increase in flexibility at H53 and N65 is much smaller than the increase observed in **Mel/Gal-10_cf_**. Moreover, Q55 and W72 in **Mel/E33A-Gal-10_cf_** have a negative *ΔB’* value, which indicates enhanced rigidity at these residues. **Raf/E33A-Gal-10_cf_** indicates a decrease in flexibility at Q55, R61, W72, and R60’ which is similar to the observations of **Mel/E33A-Gal-10_cf_** except for R61.

### All-atom MD simulation of melezitose bound Gal-10

All-atom molecular dynamics (MD) simulations using AMBER were performed to investigate the structural stabilization of melezitose in the dimer and crystal forms (See the Experimental Procedures in SI). The simulation of Gal-10 in the crystal state was evaluated using the 14-mer containing a central dimer surrounded by six dimers (Movie S1). The dimer and the 14-mer systems of **Mel/E33A-Gal-10_cf_** and **Mel/Gal-10_cf_** were subjected to 100 ns MD simulations. The retention time of melezitose was evaluated from the distance between W72 and the terminal saccharide unit, Glc^2^_Mel_ or Glc^1^_Mel_ (Figure S8a and S9a). In the dimer of **Mel/E33A-Gal-10_cf_**, one of the two melezitose molecules was retained for 100 ns and in the 14-mer, two melezitose molecules were retained over one run, while one of the two was retained in the other two runs (Figure S8b,c). These results suggest that the 14-mer retains melezitose more effectively than the dimer. In the dimer of **Mel/Gal-10_cf_**, two melezitose molecules were retained over one run, while one of the two was retained in the other runs (Figure S9b). In the 14-mer of **Mel/Gal-10_cf_**, two melezitose molecules were retained over one run, although either one or both were released in the other runs (Figure S9c). This result suggests that the dimer of **Mel/Gal-10_cf_** retains melezitose more effectively than the 14-mer, in contrast with the results observed for **Mel/E33A-Gal-10_cf_**. Furthermore, 14-mer **Mel/E33A-Gal-10_cf_** has higher efficiency than **Mel/Gal-10_cf_** in retaining melezitose.

The conformational variation of melezitose bound to Gal-10 was evaluated by observations of the dihedral angle of the glycosidic linkage, (φ1, ψ1) and (φ2, ψ2), with two melezitose molecules at the binding sites of the Gal-10 dimer at the center of the 14-mer (Figure 4a and S8a,b). The distribution of (φ1, ψ1) is localized in one area (Area 1) widely in **Mel/E33A-Gal-10_cf_**, whereas it was localized in two areas (Areas 2 and 3) in **Mel/Gal-10_cf_** (Figure 4b,c). On the other hand, the distribution of (φ2, ψ2) is localized in three areas (Areas 4, 5, and 6) in **Mel/E33A-Gal-10_cf_**, and localized in one area (Area 7) in **Mel/Gal-10_cf_** (Figure 4d,e). These results indicate that the dynamics of melezitose in the binding site of Gal-10 are different in the presence of the mutation at E33’.

**Figure 4.**
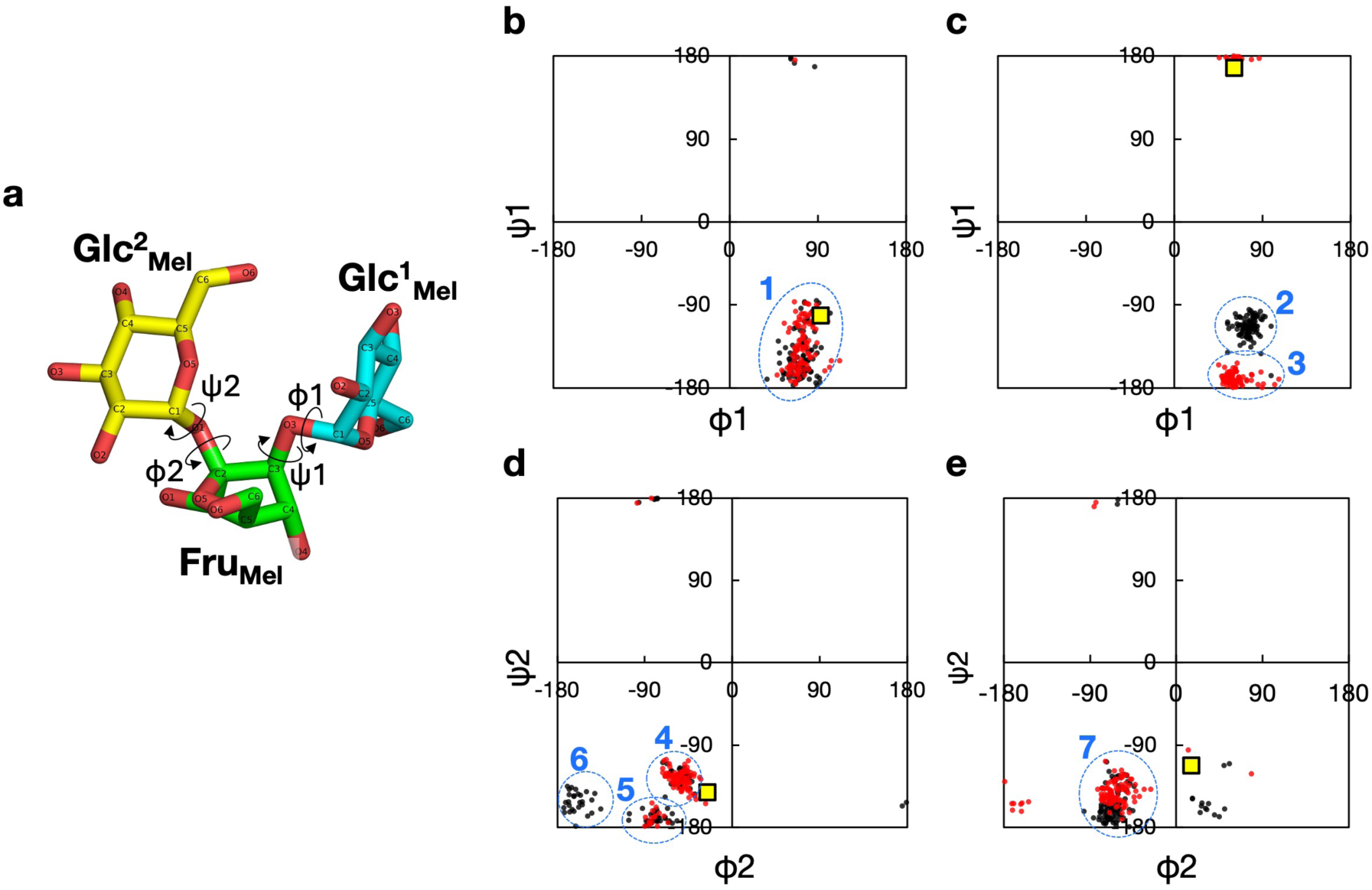
Analysis of dihedral angles of glycosidic linkages in melezitose. (a) Dihedral angles, (φ1, ψ1) and (φ2, ψ2), of melezitose. The distributions of (φ1, ψ1) in (b) **Mel/E33A-Gal-10_cf_** and (c) **Mel/Gal-10_cf_**. The distributions of (φ2, ψ2) in (d) **Mel/E33A-Gal-10_cf_** and (e) **Mel/Gal-10_cf_**. The areas where the angular distribution is localized are designated as Area 1-7 in (b-e), surrounded by a blue dotted line, respectively.

## Discussion

We determined the structures of trisaccharides and disaccharides using 50–200 µm **Gal-10_cf_** synthesized at a reaction scale of 80 µL by CFPC. The crystal structures of the complexes of saccharides and **Gal-10_cf_** determined at 1.57–2.31 Å resolution indicate the same space group and identical cell parameters as those of Gal-10_iv_. The large crystal size of **Gal-10_cf_**, which is challenging to obtain by the in vivo crystallization,^36^ enabled us to collect the diffraction data, indicating that **Gal-10_cf_** has the potential for high-throughput screening to investigate binding of small substrates.

X-ray crystal structure analyses for the crystals of complexes of trisaccharides with **Gal-10_cf_** or **E33A-Gal-10_cf_** provide the first structures of melezitose and raffinose bound to Gal-10. Raffinose is particularly interesting due to its role as a prebiotic promoting healthy gut microbiota.^37^ The structural insights gained from this study are expected to contribute to a better understanding of how raffinose interacts with biological macromolecules, including enzymes and bacterial receptors in the gut.^37^ Melezitose is an effective inhibitor for testing the affinity of the saccharide chains of yeasts for binding to lectins in the drug design process.^38^ **Mel/E33A-Gal-10_cf_** decreases the average *B*-factor of melezitose compared to **Gal-10_cf_**. Melezitose in **Mel/E33A-Gal-10_cf_** is fixed at Glc^1^_Mel_ by the hydrogen bond with R60’, which is restricted in **Mel/Gal-10_cf_** due to the steric hindrance with E33’. Additionally, the another terminal monosaccharide unit, Glc^2^_Mel_, are are captured by key amino acid residues (Figure 3a and 5). In **Mel/Gal-10_cf_**, both melezitose and surrounding residues fluctuate to retain the same number of hydrogen bonds and hydrophobic interactions within the restricted space. The spatial configuration is a factor that regulates the rigidity of non-covalent bonds between surrounding residues and trisaccharides. Earlier NMR and MD simulations of galectin and saccharides have referred to the effect of the increase of conformational entropy of galectin to maintain the binding between the saccharide and galectin.^39, 40^ Our observations indicate that residues surrounding the trisaccharide fluctuate to a greater extent relative to the unbound state in **Mel/Gal-10_cf_**, which suggests that the increase of conformational entropy contributes favorably to saccharide binding. In **Mel/E33A-Gal-10_cf_**, the surrounding residues exhibit a loss of conformational entropy upon binding. However, the entropy loss is sufficiently small to be compensated by hydrogen bonds and hydrophobic interactions due to the rigid protein structure in its unbound state (Figure 2a,c). The results derived from the *ΔB’* analysis provide experimental support for the theoretical analysis conducted by NMR and MD simulations.

**Figure 5.**
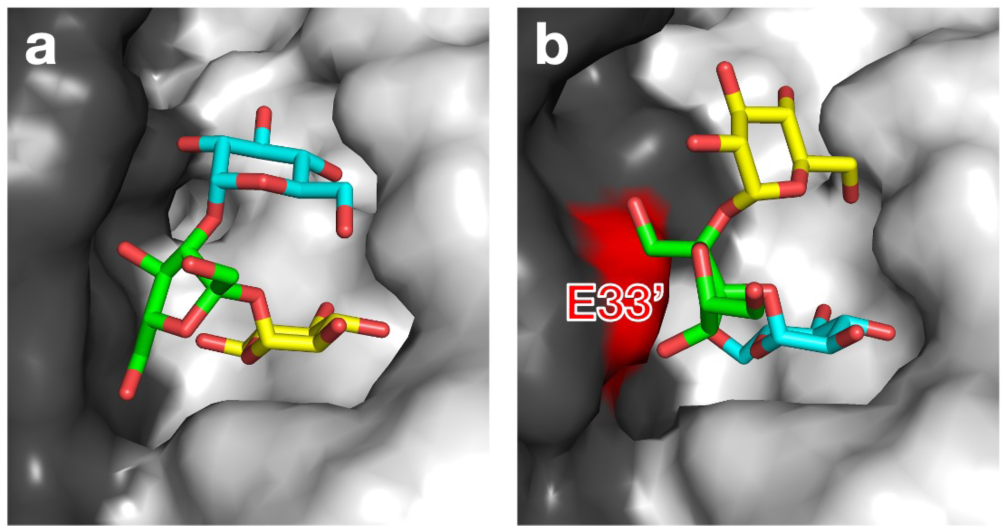
The spatial configuration of the binding site of (a) **Mel/E33A-Gal-10_cf_** and (b) **Mel/Gal-10_cf_**. The stick models show the melezitose. The Glc^1^_Mel_, Fru_Mel_, and Glc^2^_Mel_ in melezitose are colored cyan, green, and yellow, respectively. The surface model shows the Gal-10, in which each monomer is colored gray or black.

MD simulations of **Mel/E33A-Gal-10_cf_** and **Mel/Gal-10_cf_** revealed that the binding site restricts the flexibility of the glycosidic linkage in melezitose. The distribution of inner dihedral angles of melezitose in **Mel/E33A-Gal-10_cf_** and **Mel/Gal-10_cf_** is localized to two or three areas, respectively (Figure 4). Because the conformation of saccharide contributes to the protein binding affinity, the molecular dynamics of the saccharide molecule is an important factor in understanding the bio-signaling which occurs via binding of saccharide and this information will be used to design inhibitors for drug discovery. For example, the MD simulation of mannose-bound cyanoviron, an anti-HIV virucidal protein, indicates that the dihedral angle of glycosidic linkage in mannose is restricted in the high-affinity binding site rather than in the low-affinity site.^41^ Investigations of the affinity of the saccharide in the presence of specific protein mutations introduced in MD simulations of the crystal structure obtained by CFPC will enable us to characterize the relationship between the saccharide dynamics and the binding affinity of the sugar-binding protein.

## Conclusion

In conclusion, we demonstrated a rapid and purification-free method for producing high-quality Gal-10 crystals through cell-free protein crystallization (CFPC), enabling atomic-level structural analysis of flexible disaccharides and trisaccharides. The successful determination of the previously unreported structures of melezitose and raffinose highlights the potential of this method for resolving glycan conformations that are otherwise difficult to capture. By leveraging engineered variants such as E33A, we demonstrated that subtle modifications to the binding pocket can significantly enhance saccharide immobilization by modulating hydrogen bonding networks and conformational dynamics. These insights are supported by the results of MD simulations and *ΔB’* analyses, providing a quantitative framework to dissect the entropy-enthalpy compensation in saccharide recognition. This study establishes Gal-10 as an adaptable scaffold for glycan structure analysis and sets the foundation for creating tailored crystal libraries that accommodate a broad range of carbohydrate ligands. Future directions include integrating this platform with high-throughput screening systems and computational design strategies to accelerate glycan-targeted drug discovery, prebiotic functional screening, and rational design of lectin inhibitors. Ultimately, this strategy offers a powerful toolkit for probing glycan-mediated biological processes with unprecedented structural and dynamic insights.

## ASSOCIATED CONTENT

### Supporting Information

The following files are available free of charge.

Detailed experimental procedures for cell-free protein synthesis and crystallization, MALDI-TOF/MS, adsorption of tri- and disaccharides into protein crystal, X-ray crystal structure analysis, analysis of normalized *B*-factor, and molecular dynamics simulation, detailed data of structural analysis and molecular dynamics simulation (PDF)

**Movie S1.** Molecular dynamics simulation for 14-mer of **Mel/E33A-Gal-10_cf_** (AVI)

### Author Contributions

The manuscript was written through contributions of all authors. M.K., S.A., and T.U. designed research; X.Y., R.K., T.S., and M.K. performed research; M.K., T.F., and K.H. analyzed data; and M.K., S.A., and T.U. wrote the paper. All authors have approved the final version of the manuscript.

### Notes

The authors declare no competing financial interest.

## ACKNOWLEDGMENT

This work was supported by JSPS KAKENHI Grant No. JP22H00347, JP25H02254 to T.U. and JP22K19266 to S.A., and the Adaptable and Seamless Technology Transfer Program through Target-driven R&D (JPMJTR20U1, JPMJTR224A) from the Japan Science and Technology Agency to T.U. Synchrotron radiation experiments were conducted under the approval of 2021A2772, 2021B2772, 2021A2744, 2021B2744, 2022A2735, 2022B2735, 2022A2771, and 2022B2771, 2023A2745, and 2023B2745 at SPring-8. This work was supported by SUNBOR SCHOLARSHIP from the Suntory Foundation for Life Sciences to M.K. This research was partially supported by the Platform Project for Supporting Drug Discovery and Life Science Research [Basis for Supporting Innovative Drug Discovery and Life Science Research (BINDS)] from the AMED under Grant number JP21am0101070 (support number 1854).

## ABBREVIATIONS

Gal-10: galectin-10;
CFPC: cell-free protein crystallization;
*B′*: Normalized *B*-factor;
IDPs: intrinsically disordered proteins;
RMSD: root-mean-square deviations.

## For Table of Contents Only

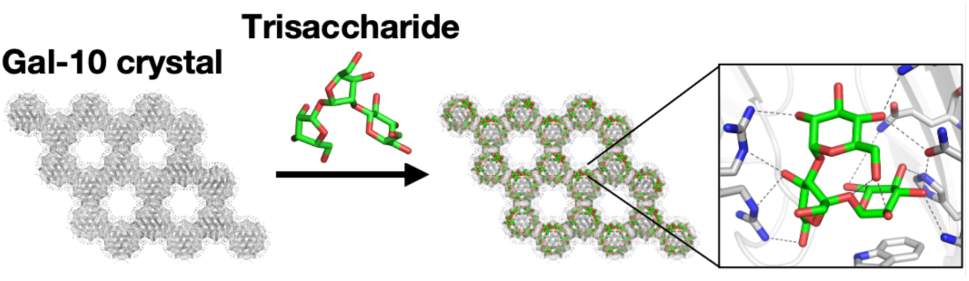

The structure of trisaccharide was determined using Galectin-10 (Gal-10) crystals prepared by cell-free protein crystallization, offering insights into the dynamics of saccharide in the binding site.

## REFERENCES

(1) de Toro, B. F.; Peng, W. J.; Thompson, A. J.; Domínguez, G.; Cañada, F. J.; Pérez-Castells, J.; Paulson, J. C.; Jiménez-Barbero, J.; Canales, A. Avenues to Characterize the Interactions of Extended N-Glycans with Proteins by NMR Spectroscopy: The Influenza Hemagglutinin Case. Angew. Chem. Int. Ed. 2018, 57 (46), 15051–15055. DOI: 10.1002/anie.201807162.

(2) Nagae, M.; Kanagawa, M.; Morita-Matsumoto, K.; Hanashima, S.; Kizuka, Y.; Taniguchi, N.; Yamaguchi, Y. Atomic visualization of a flipped-back conformation of bisected glycans bound to specific lectins. Sci. Rep. 2016, 6. DOI: 10.1038/srep22973.

(3) Canales, A.; Mallagaray, A.; Pérez-Castells, J.; Boos, I.; Unverzagt, C.; André, S.; Gabius, H. J.; Cañada, F. J.; Jiménez-Barbero, J. Breaking Pseudo-Symmetry in Multiantennary Complex *N*-Glycans Using Lanthanide-Binding Tags and NMR Pseudo-Contact Shifts. Angew. Chem. Int. Ed. 2013, 52 (51), 13789–13793. DOI: 10.1002/anie.201307845.

(4) Canales, A.; Boos, I.; Perkams, L.; Karst, L.; Luber, T.; Karagiannis, T.; Domínguez, G.; Cañada, F. J.; Pérez-Castells, J.; Häussinger, D.;, et al. Breaking the Limits in Analyzing Carbohydrate Recognition by NMR Spectroscopy: Resolving Branch-Selective Interaction of a Tetra-Antennary *N*-Glycan with Lectins. Angew. Chem. Int. Ed. 2017, 56 (47), 14987–14991. DOI: 10.1002/anie.201709130.

(5) Gimeno, A.; Valverde, P.; Ardá, A.; Jiménez-Barbero, J. Glycan structures and their interactions with proteins. A NMR view. Curr. Opin. Struct. Biol. 2020, 62, 22–30. DOI: 10.1016/j.sbi.2019.11.004.

(6) Wormald, M. R.; Petrescu, A. J.; Pao, Y. L.; Glithero, A.; Elliott, T.; Dwek, R. A. Conformational studies of oligosaccharides and glycopeptides: Complementarity of NMR, X-ray crystallography, and molecular modelling. Chem. Rev. 2002, 102 (2), 371–386. DOI: 10.1021/cr990368i.

(7) Woods, R. J. Predicting the Structures of Glycans, Glycoproteins, and Their Complexes. Chem. Rev. 2018, 118 (17), 8005–8024. DOI: 10.1021/acs.chemrev.8b00032.

(8) Yu, Y.; Delbianco, M. Conformational Studies of Oligosaccharides. Chem. Eur. J. 2020, 26 (44), 9814–9825. DOI: 10.1002/chem.202001370.

(9) Meredith, R. J.; Carmichael, I.; Woods, R. J.; Serianni, A. S. *MA’AT* Analysis: Probability Distributions of Molecular Torsion Angles in Solution from NMR Spectroscopy. Acc. Chem. Res. 2023, 56 (17), 2313–2328. DOI: 10.1021/acs.accounts.3c00286.

(10) Widmalm, G. Glycan Shape, Motions, and Interactions Explored by NMR Spectroscopy. Jacs Au 2024, 4 (1), 20–39. DOI: 10.1021/jacsau.3c00639.

(11) Pearson, A. D.; Mills, J. H.; Song, Y. F.; Nasertorabi, F.; Han, G. W.; Baker, D.; Stevens, R. C.; Schultz, P. G. Trapping a transition state in a computationally designed protein bottle. Science 2015, 347 (6224), 863–867. DOI: 10.1126/science.aaa2424.

(12) Yamasaki, S.; Nikaido, E.; Nakashima, R.; Sakurai, K.; Fujiwara, D.; Fujii, I.; Nishino, K. The crystal structure of multidrug-resistance regulator RamR with multiple drugs. Nat. Commun. 2013, 4, 2078. DOI: 10.1038/ncomms3078.

(13) Yamasaki, S.; Nakashima, R.; Sakurai, K.; Baucheron, S.; Giraud, E.; Doublet, B.; Cloeckaert, A.; Nishino, K. Crystal structure of the multidrug resistance regulator RamR complexed with bile acids. Sci. Rep. 2019, 9, 177. DOI: 10.1038/s41598-018-36025-8.

(14) Maita, N. Crystal structure determination of ubiquitin by fusion to a protein that forms a highly porous crystal lattice. J. Am. Chem. Soc. 2018, 140 (42), 13546–13549. DOI: 10.1021/jacs.8b07512.

(15) Kojima, M.; Abe, S.; Furuta, T.; Tran, D. P.; Hirata, K.; Yamashita, K.; Hishikawa, Y.; Kitao, A.; Ueno, T. Engineering of an in-cell protein crystal for fastening a metastable conformation of a target miniprotein. Biomater. Sci. 2023, 11 (4), 1350–1357. DOI: 10.1039/d2bm01759h.

(16) Abe, S.; Tanaka, J.; Kojima, M.; Kanamaru, S.; Hirata, K.; Yamashita, K.; Kobayashi, A.; Ueno, T. Cell-free protein crystallization for nanocrystal structure determination. Sci. Rep. 2022, 12 (1), 16031. DOI: 10.1038/s41598-022-19681-9.

(17) Kojima, M.; Abe, S.; Furuta, T.; Hirata, K.; Yao, X.; Kobayashi, A.; Kobayashi, R.; Ueno, T. High-throughput structure determination of an intrinsically disordered protein using cell-free protein crystallization. Proc. Natl. Acad. Sci. U.S.A. 2024, 121 (25). DOI: 10.1073/pnas.2322452121.

(18) Dyer, K. D.; Rosenberg, H. F. Eosinophil Charcot-Leyden crystal protein binds to beta-galactoside sugars. Life Sci. 1996, 58 (23), 2073–2082. DOI: 10.1016/0024-3205(96)00201-9.

(19) Su, J. Y. A Brief History of Charcot-Leyden Crystal Protein/Galectin-10 Research. Molecules 2018, 23 (11), 2931. DOI: 10.3390/molecules23112931.

(20) Allen, J. E.; Sutherland, T. E. Crystal-clear treatment for allergic disease. Science 2019, 364 (6442), 738–739. DOI: 10.1126/science.aax6175.

(21) Ueki, S.; Miyabe, Y.; Yohei, Y.; Fukuchi, M.; Hirokawa, M.; Spencer, L. A.; Weller, P. F. Charcot-Leyden Crystals in Eosinophilic Inflammation: Active Cytolysis Leads to Crystal Formation. Curr. Allergy Asthma Rep. 2019, 19 (8), 35. DOI: 10.1007/s11882-019-0868-0.

(22) Swaminathan, G. J.; Leonidas, D. D.; Savage, M. P.; Ackerman, S. J.; Acharya, K. R. Selective recognition of mannose by the human eosinophil Charcot-Leyden crystal protein (galectin-10): A crystallographic study at 1.8 angstrom resolution. Biochemistry 1999, 38 (42), 13837–13843. DOI: 10.1021/bi990756e.

(23) Ackerman, S. J.; Liu, L.; Kwatia, M. A.; Savage, M. P.; Leonidas, D. D.; Swaminathan, G. J.; Acharya, K. R. Charcot-Leyden crystal protein (galectin-10) is not a dual function galectin with lysophospholipase activity but binds a lysophospholipase inhibitor in a novel structural fashion. J. Biol. Chem. 2002, 277 (17), 14859–14868. DOI: 10.1074/jbc.M200221200.

(24) Su, J. Y.; Gao, J.; Si, Y. L.; Cui, L. L.; Song, C. Y.; Wang, Y.; Wu, R. J.; Tai, G. H.; Zhou, Y. F. Galectin-10: a new structural type of prototype galectin dimer and effects on saccharide ligand binding. Glycobiology 2018, 28 (3), 159–168. DOI: 10.1093/glycob/cwx107.

(25) Persson, E. K.; Verstraete, K.; Heyndrickx, I.; Gevaert, E.; Aegerter, H.; Percier, J. M.; Deswarte, K.; Verschueren, K. H. G.; Dansercoer, A.; Gras, D.;, et al. Protein crystallization promotes type 2 immunity and is reversible by antibody treatment. Science 2019, 364 (6442), eaaw4295. DOI: 10.1126/science.aaw4295.

(26) Su, J. Y.; Song, C. Y.; Si, Y. L.; Cui, L. L.; Yang, T.; Li, Y. Y.; Wang, H.; Tai, G. H.; Zhou, Y. F. Identification of key amino acid residues determining ligand binding specificity, homodimerization and cellular distribution of human galectin-10. Glycobiology 2019, 29 (1), 85–93. DOI: 10.1093/glycob/cwy087.

(27) Itoh, A.; Nonaka, Y.; Nakakita, S. I.; Yoshida, H.; Nishi, N.; Nakamura, T.; Kamitori, S. Structures of human galectin-10/monosaccharide complexes demonstrate potential of monosaccharides as effectors in forming Charcot-Leyden crystals. Biochem. Biophys. Res. Commun. 2020, 525 (1), 87–93. DOI: 10.1016/j.bbrc.2020.02.037.

(28) Kaneko, T.; Kohmoto, T.; Kikuchi, H.; Shiota, M.; Iino, H.; Mitsuoka, T. EFFECTS OF ISOMALTOOLIGOSACCHARIDES WITH DIFFERENT DEGREES OF POLYMERIZATION ON HUMAN FECAL BIFIDOBACTERIA. *Biosci., Biotechnol.*, Biochem. 1994, 58 (12), 2288–2290. DOI: 10.1271/bbb.58.2288.

(29) Kaplan, H.; Hutkins, R. W. Fermentation of fructooligosaccharides by lactic acid bacteria and bifidobacteria. Appl. Environ. Microbiol. 2000, 66 (6), 2682–2684. DOI: 10.1128/aem.66.6.2682-2684.2000.

(30) Sanz, M. L.; Côté, G. L.; Gibson, G. R.; Rastall, R. A. Prebiotic properties of alternansucrase maltose-acceptor oligosaccharides. J. Agric. Food. Chem. 2005, 53 (15), 5911–5916. DOI: 10.1021/jf050344e.

(31) Martínez-Villaluenga, C.; Cardelle-Cobas, A.; Olano, A.; Corzo, N.; Villamiel, M.; Jimeno, M. L. Enzymatic synthesis and identification of two trisaccharides produced from lactulose by transgalactosylation. J. Agric. Food. Chem. 2008, 56 (2), 557–563. DOI: 10.1021/jf0721343.

(32) Fushinobu, S.; Abou Hachem, M. Structure and evolution of the bifidobacterial carbohydrate metabolism proteins and enzymes. Biochem. Soc. Trans. 2021, 49 (2), 563–578. DOI: 10.1042/bst20200163.

(33) Sawasaki, T.; Ogasawara, T.; Morishita, R.; Endo, Y. A cell-free protein synthesis system for high-throughput proteomics. Proc. Natl. Acad. Sci. U.S.A. 2002, 99 (23), 14652–14657. DOI: 10.1073/pnas.232580399.

(34) Baker, E. N.; Hubbard, R. E. Hydrogen bonding in globular proteins. Prog. Biophys. Mol. Biol. 1984, 44 (2), 97–179. DOI: 10.1016/0079-6107(84)90007-5.

(35) Johnson, T. W.; Gallego, R. A.; Brooun, A.; Gehlhaar, D.; McTigue, M. Reviving B-Factors: Retrospective Normalized B-Factor Analysis of c-ros Oncogene 1 Receptor Tyrosine Kinase and Anaplastic Lymphoma Kinase L1196M with Crizotinib and Lorlatinib. ACS Med. Chem. Lett. 2018, 9 (9), 878–883. DOI: 10.1021/acsmedchemlett.8b00147.

(36) Ayres, W. W. PRODUCTION OF CHARCOT-LEYDEN CRYSTALS FROM EOSINOPHILS WITH AEROSOL-MA. Blood 1949, 4 (5), 595–602.

(37) Kanwal, F.; Ren, D. X.; Kanwal, W.; Ding, M. Y.; Su, J. Q.; Shang, X. Y. The potential role of nondigestible Raffinose family oligosaccharides as prebiotics. Glycobiology 2023, 33 (4), 274–288. DOI: 10.1093/glycob/cwad015.

(38) Ghazarian, A.; Oppenheimer, S. B. Microbead analysis of cell binding to immobilized lectin. Part II: Quantitative kinetic profile assay for possible identification of anti-infectivity and anti-cancer reagents. Acta Histochem. 2014, 116 (8), 1514–1518. DOI: 10.1016/j.acthis.2014.07.015.

(39) Nesmelova, I. V.; Ermakova, E.; Daragan, V. A.; Pang, M.; Menéndez, M.; Lagartera, L.; Solís, D.; Baum, L. G.; Mayo, K. H. Lactose Binding to Galectin-1 Modulates Structural Dynamics, Increases Conformational Entropy, and Occurs with Apparent Negative Cooperativity. J. Mol. Biol. 2010, 397 (5), 1209–1230. DOI: 10.1016/j.jmb.2010.02.033.

(40) Guardia, C. M. A.; Gauto, D. F.; Di Lella, S.; Rabinovich, G. A.; Martí, M. A.; Estrin, D. A. An Integrated Computational Analysis of the Structure, Dynamics, and Ligand Binding Interactions of the Human Galectin Network. J. Chem. Inf. Model. 2011, 51 (8), 1918–1930. DOI: 10.1021/ci200180h.

(41) Margulis, C. J. Computational study of the dynamics of mannose disaccharides free in solution and bound to the potent anti-HIV virucidal protein cyanovirin. J. Phys. Chem. B 2005, 109 (8), 3639–3647. DOI: 10.1021/jp0406971.

