## Supporting information for "Cell-Free Protein Crystallization Enables Rapid Structure Determination of Disaccharides and Trisaccharides Using Galectin-10 Crystals"

### Table of Contents

**Experimental Procedures**
Materials 3
Cell-free synthesis and crystallization of **Gal-10$_{cf}$** 3
MALDI-TOF MS 3
Adsorption of tri- and disaccharides into **Gal-10$_{cf}$** 3
X-ray crystal structure analysis 3
Analysis of normalized *B*-factor 4
Molecular dynamics simulation 4
**Figure S1** 6
**Figure S2** 7
**Figure S3** 8
**Figure S4** 9
**Figure S5** 10
**Figure S6** 11
**Figure S7** 12
**Figure S8** 13
**Figure S9** 14
**Table S1** 15
**References** 16

### Experimental Procedures

#### Materials

The competent cells of DH5α *E. coli* were purchased from Toyobo and InvitrogenTM (Life Technologies). The PCR reactions were implemented using QuikChange® Site-Directed Mutagenesis Kit (Stratagene) and KOD Plus Mutagenesis Kit (Toyobo). Other reagents were purchased from TCI, Wako, Nacalai Tesque, Sigma–Aldrich, and Life Technologies and were used without further purification.

#### Cell-free synthesis and crystallization of Gal-10$_{cf}$

Expression and crystallization of Gal-10 crystals were performed using a WEPRO7240 Expression Kit (CellFree Sciences). The gene of Gal-10 was cloned into the PEU-E01-MCS vector (CellFree Sciences) for the Gal-10 expression. The plasmid was amplified in DH5α bacteria and purified using the Qiagen Plasmid Midi Kit. Transcription was performed by dialysis system according to the protocol of the expression kit. After incubation for 6 h at 37 °C, mRNA was used for translation. Translation reactions were carried out using the dialysis method. 80 μL of reaction mixture containing 20 μL of WEPRO7240 wheat germ extract, 20 μL of prepared mRNA, 40 μL of SUB-AMIX SGC solution, and 40 mg/mL creatine kinase was dialyzed against 2.5 mL of SUB-AMIX SGC solution in a cylindrical cup and incubated at 20 °C for 72 h. The crystals were collected by centrifuge and washed with PBS several times.

#### MALDI-TOF MS

The identities of **Gal-10$_{cf}$** were confirmed by MALDI-TOF-MS using a Bruker ultrafleXtreme. The molecular weight of the sequence lacking the N-terminal methionine was calculated as the theoretical value.

#### Adsorption of tri- and disaccharides into Gal-10$_{cf}$

Gal-10 crystals were soaked in a 200 μL aqueous solution of melezitose (1.0 M), raffinose (the saturated concentration, ca. 0.4 M), sucrose (1.0 M), trehalose (1.0 M), maltose (1.0 M), lactose (the saturated concentration, ca. 0.6 M) at 20 °C for 24 h to adsorb the saccharide molecules into Gal-10 crystals.

#### X-ray crystal structure analysis

Before the data collection, the crystals were immersed in PBS buffer containing 50% ethylene glycol and then loaded onto the MicroLoops (Mitegen), followed by freezing in liquid nitrogen. The composite crystals incubated with di- and trisaccharides were immersed in cryoprotectant (50% ethylene glycol) containing di- and trisaccharides under the same condition for soaking for X-ray structure determination. X-ray diffraction data of **Gal-10$_{cf}$**, **Mel/Gal-10$_{cf}$**, **Raf/Gal-10$_{cf}$**, **Suc/Gal-10$_{cf}$**, **Tre/Gal-10$_{cf}$**, **Mal/Gal-10$_{cf}$**, **Lac/Gal-10$_{cf}$**, **Mel/E33A-Gal-10$_{cf}$**, and **Raf/E33A-Gal-10$_{cf}$**, were collected at 100 K at beamline BL32XU at SPring-8 using X-ray wavelength of 1.00 Å. The complete sets of structure factor amplitudes were obtained by merging multiple small-wedge (5° or 10° each) datasets collected from single crystals. The crystal positions in a cryoloop were identified by low-dose raster scan. The whole data collection process was automated by ZOO system including sample exchange by a robot. Collected datasets were automatically processed and merged by KAMO.[1] Each dataset was indexed and integrated using XDS.[2] The datasets consistently indexed with the known cell parameter were selected and two possible reindex operators (hkl and -hlk) were tested to give better match to the previously solved data (1LCL). The datasets were subjected to hierarchical clustering by pairwise correlation coefficient of intensities. The datasets in each cluster were scaled and merged using XSCALE[2] with outlier rejections implemented in KAMO.[2] The clusters with the highest CC1/2 were chosen for downstream analyses. The structure was solved by rigid body refinement with phenix.refine using the previously solved structure (PDB ID: 1LCL).[3] Refinement of the protein structure was performed at resolutions of 1.60 Å for **Gal-10$_{cf}$**, 1.99 Å for **Mel/Gal-10$_{cf}$**, 1.81 Å for **Raf/Gal-10$_{cf}$**, 1.57 Å for **Suc/Gal-10$_{cf}$**, 1.78 Å for **Tre/Gal-10$_{cf}$**, 2.31 Å for **Mal/Gal-10$_{cf}$**, 1.57 Å for **Lac/Gal-10$_{cf}$**, 1.62 Å for **Mel/E33A-Gal-10$_{cf}$**, 1.57 Å for **Raf/E33A-Gal-10$_{cf}$**, using REFMAC5 in the of CCP4 suite.[3] Rebuilding was performed using COOT based on sigma-A weighted $2|F_o|-|F_c|$ and $|F_o|-|F_c|$ electron density maps. The models were subjected to quality analysis during the various refinement stage with omit maps and RAMPAGE.[4] The diffraction and refinement statistics are summarized in Table S1. All images were produced using PyMOL (https://pymol.org/).

**Analysis of normalized *B*-factor**

The flexibility of the residues was evaluated by calculating the normalized *B*-factor (*B'*), which is an expression of the *B*-factor (*B*) in units of standard deviation ($\sigma B$) about the mean value ($\mu B$, eq 1).[5] Their *B'* values were individually compared with the *B'* of **Gal-10$_{cf}$** ($B'_{apo}$).[5]

$$B' = (B-\mu B)/\sigma B \tag{1}$$

$$\Delta B' = B'-B'_{apo} \tag{2}$$

**Molecular dynamics simulation**

The initial structures of dimer and 14-mer of **Mel/Gal-10$_{cf}$** and **Mel/E33A-Gal-10$_{cf}$** were constructed from the crystal structure of **Mel/Gal-10$_{cf}$** (PDB ID: 9UWN) and **Mel/E33A-Gal-10$_{cf}$** (PDB ID: 8JAE) (Movie S1). Counter ions ($Na^+$ and $Cl^-$) were added to preserve electric neutrality. For each system, the energy minimization of 300 steps was carried out with restraints of heavy atoms. Then, 500 ps equilibration under NVT condition (300 K) and 500 ps equilibration under NPT condition (300 K and 1 bar) were conducted with the same restraints above. Finally, 100 ns production runs were conducted for all systems. For 14-mer system, six dimers surrounding the center dimer in the 14-mer system were restrained by 50 kcal/mol·Å$^2$ to represent the crystal state. All the MD simulations were performed using the Amber ff19SB force field, GLYCAM force field, and TIP3P water model.[6-8] The temperature and pressure were regulated by the Berendsen thermostat and barostat, respectively. The time step was set to 2 fs, and the trajectories were recorded every 1 ns. The MDTraj software was used to analyze the distances and dihedral angles.[9]

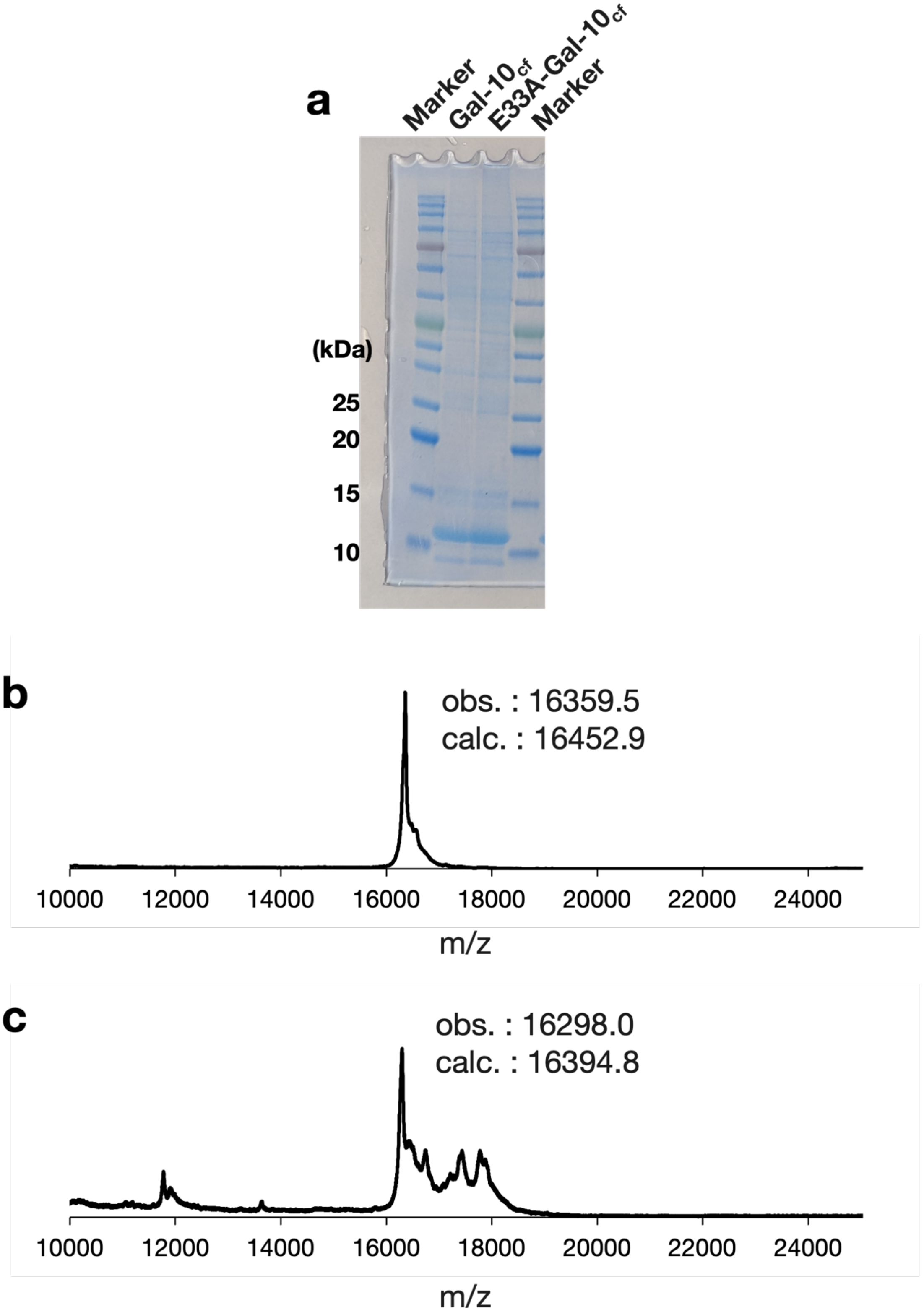


**Fig. S1** Characterization of **Gal-10$_{cf}$** by (a) SDS-PAGE, and MALDI-TOF MS of (b) **Gal-10$_{cf}$** and (c) **E33A-Gal-10$_{cf}$**. The difference in m/z values between obs. and calc. is accounted for by the loss of the N-terminal methionine and the association of a potassium ion.

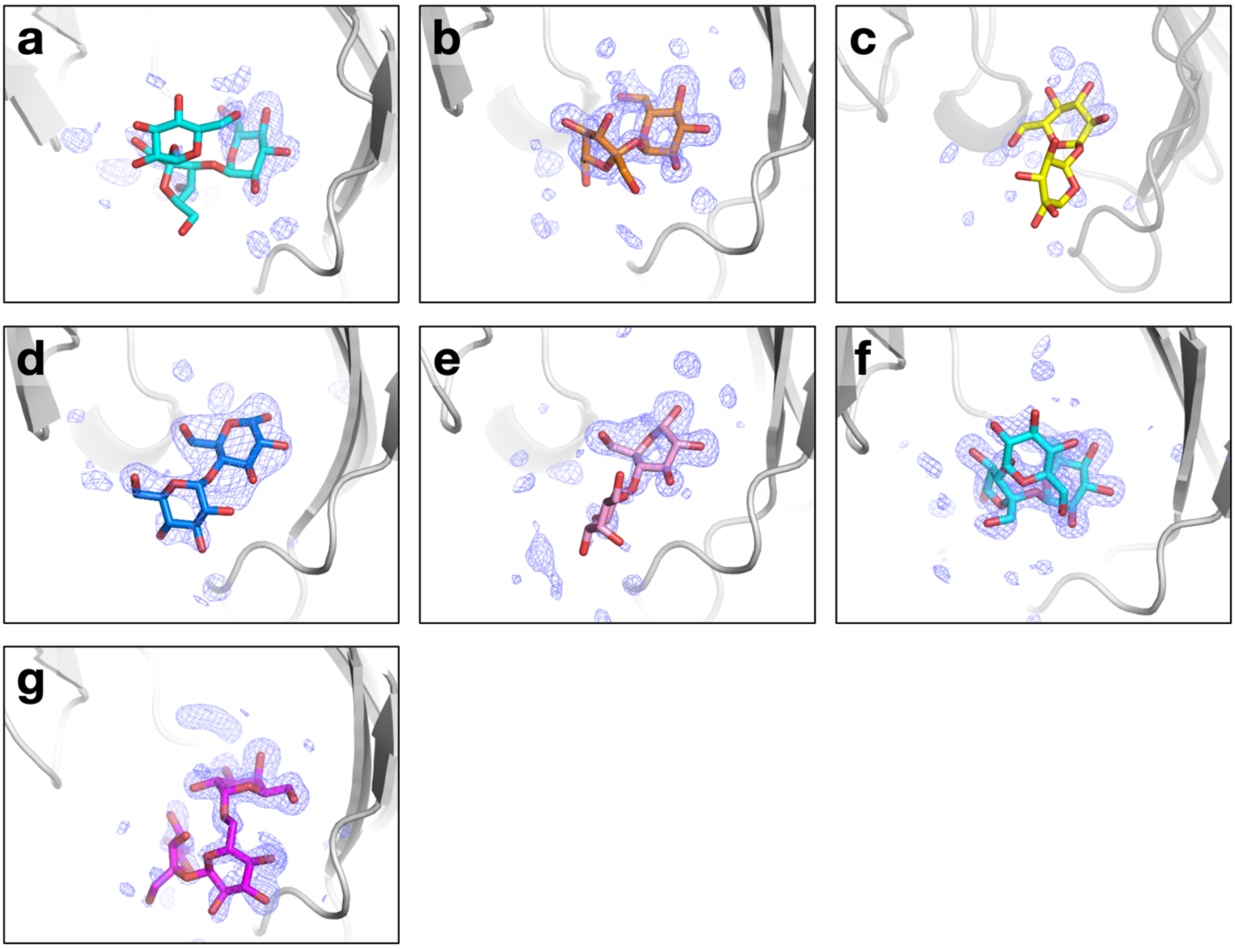


**Figure S2** Crystal structures of the saccharide-bound **Gal-10$_{cf}$s**. The polder map[10] for (a) melezitose in **Mel/Gal-10$_{cf}$**, (b) sucrose in **Suc/Gal-10$_{cf}$**, (c) trehalose in **Tre/Gal-10$_{cf}$**, (d) maltose in **Mal/Gal-10$_{cf}$**, (e) lactose in **Lac/Gal-10$_{cf}$**, (f) melezitose in **Mel/E33A-Gal-10$_{cf}$**, (g) raffinose in **Raf/E33A-Gal-10$_{cf}$**. The polder map contoured at 3.0 σ are shown in blue. Saccharides are shown as stick models. Proteins are shown as ribbon models.

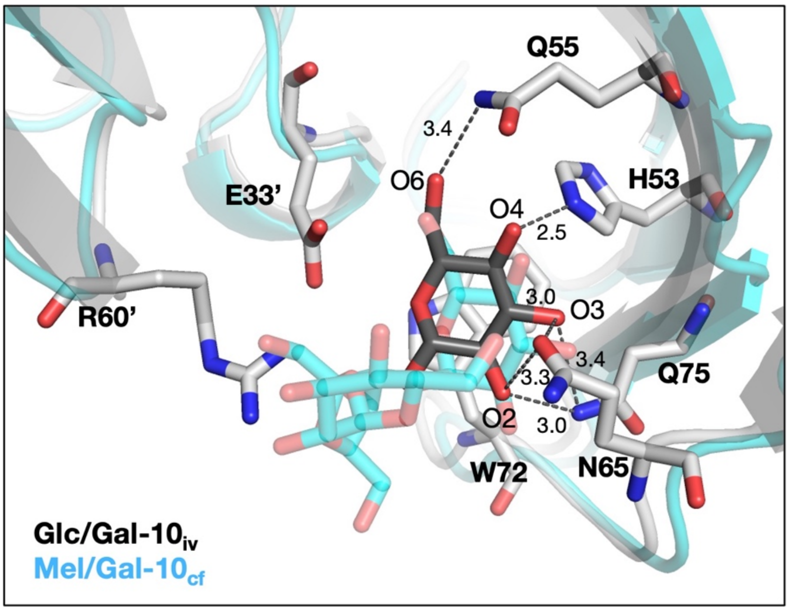


**Figure S3** Crystal structure of the glucose-bound Gal-10 (Glc/Gal-$10_{iv}$, gray) (PDB ID: 6L64) overlayed with that of **Mel/Gal-$\mathbf{10_{cf}}$** (cyan). Saccharides are shown as stick models. The structure of proteins is shown as ribbon and stick models. The cut-off distance of OH---O and NH---O hydrogen bonds is 3.5 Å.[11]

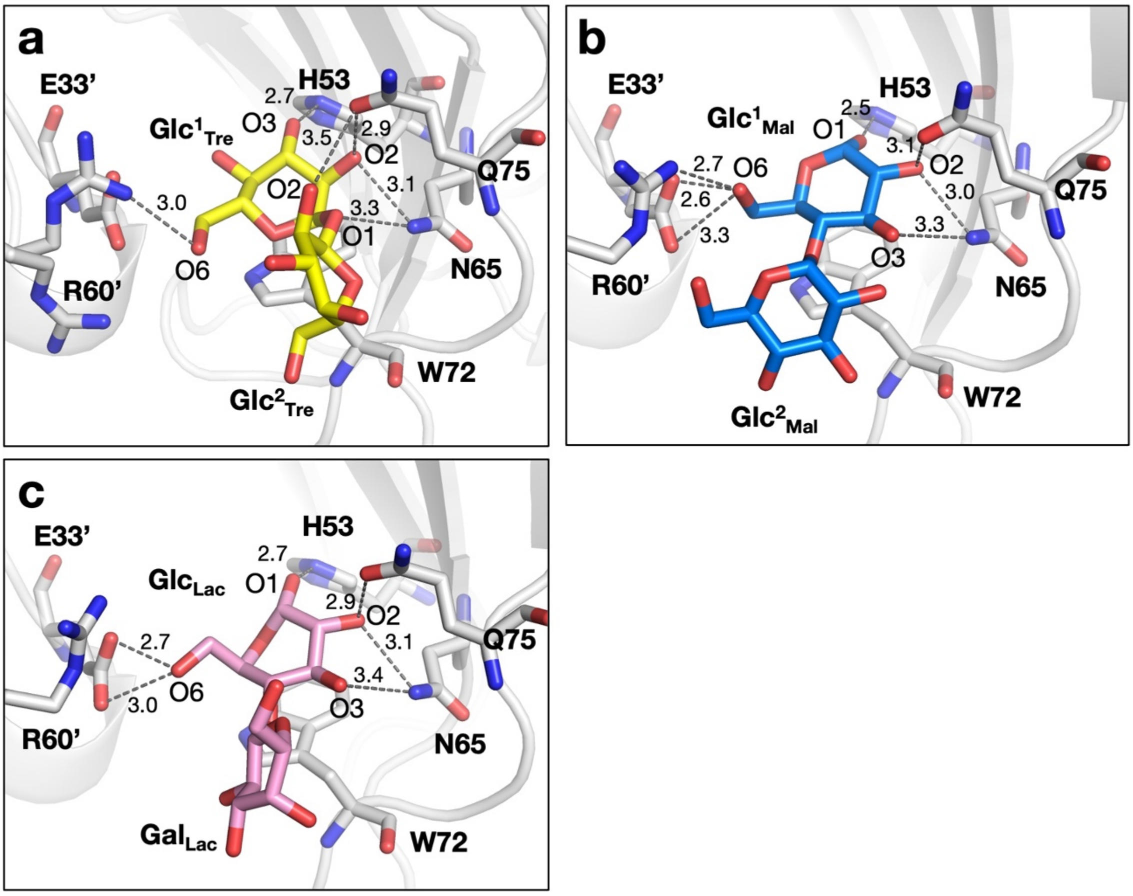


**Figure S4** Crystal structures and the noncovalent interaction networks of (a) trehalose, (b) maltose, and (c) lactose immobilized in **Gal-10$_{cf}$**. Disaccharides are shown as stick models. The structure of proteins is shown as gray-colored ribbon and stick models. The cut-off distance of OH---O and NH---O hydrogen bonds is 3.5 Å.[11]

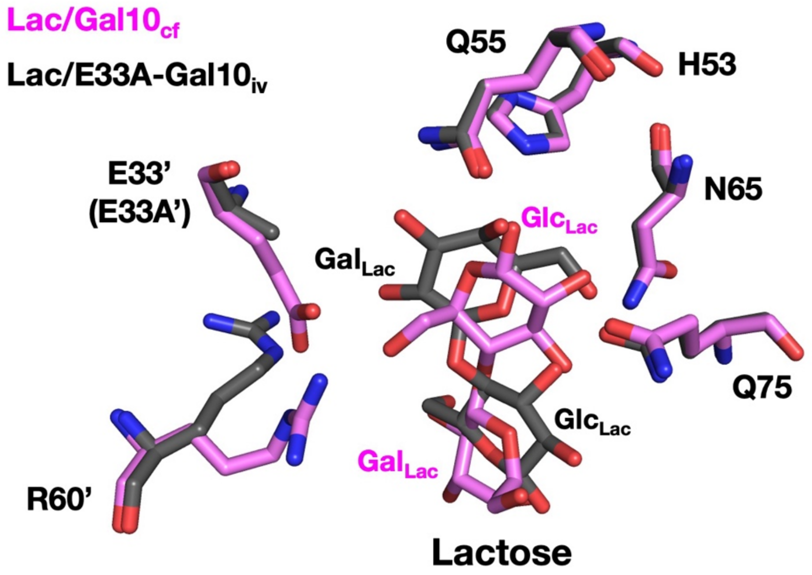


**Figure S5** The overlayed structure of lactose binding sites of **Lac/Gal-10$_{\mathbf{cf}}$** (pink) and Lac/E33A-Gal-10$_{iv}$ (PDB ID: 6A1T, black). The structure of all amino acids and disaccharides are shown as stick models.

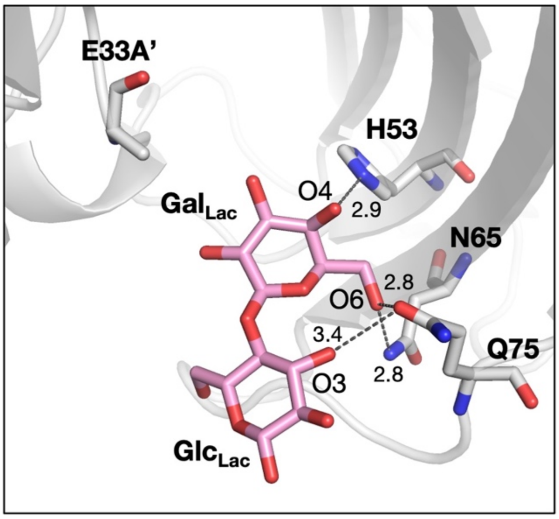


**Figure S6** Crystal structure and the noncovalent interaction networks of Lac/E33A-Gal-$10_{iv}$ (PDB ID: 6A1T).

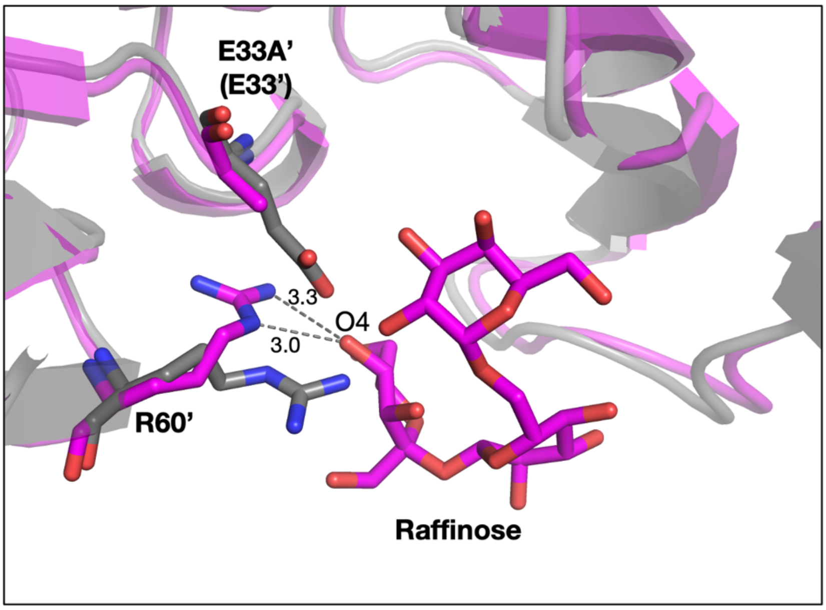


**Figure S7** The overlayed structure of raffinose binding sites of **Raf/E33A-Gal-10$_{cf}$** (magenta) and **Raf/Gal-10$_{cf}$** (gray). The structures of all amino acids and raffinose are shown as stick models.

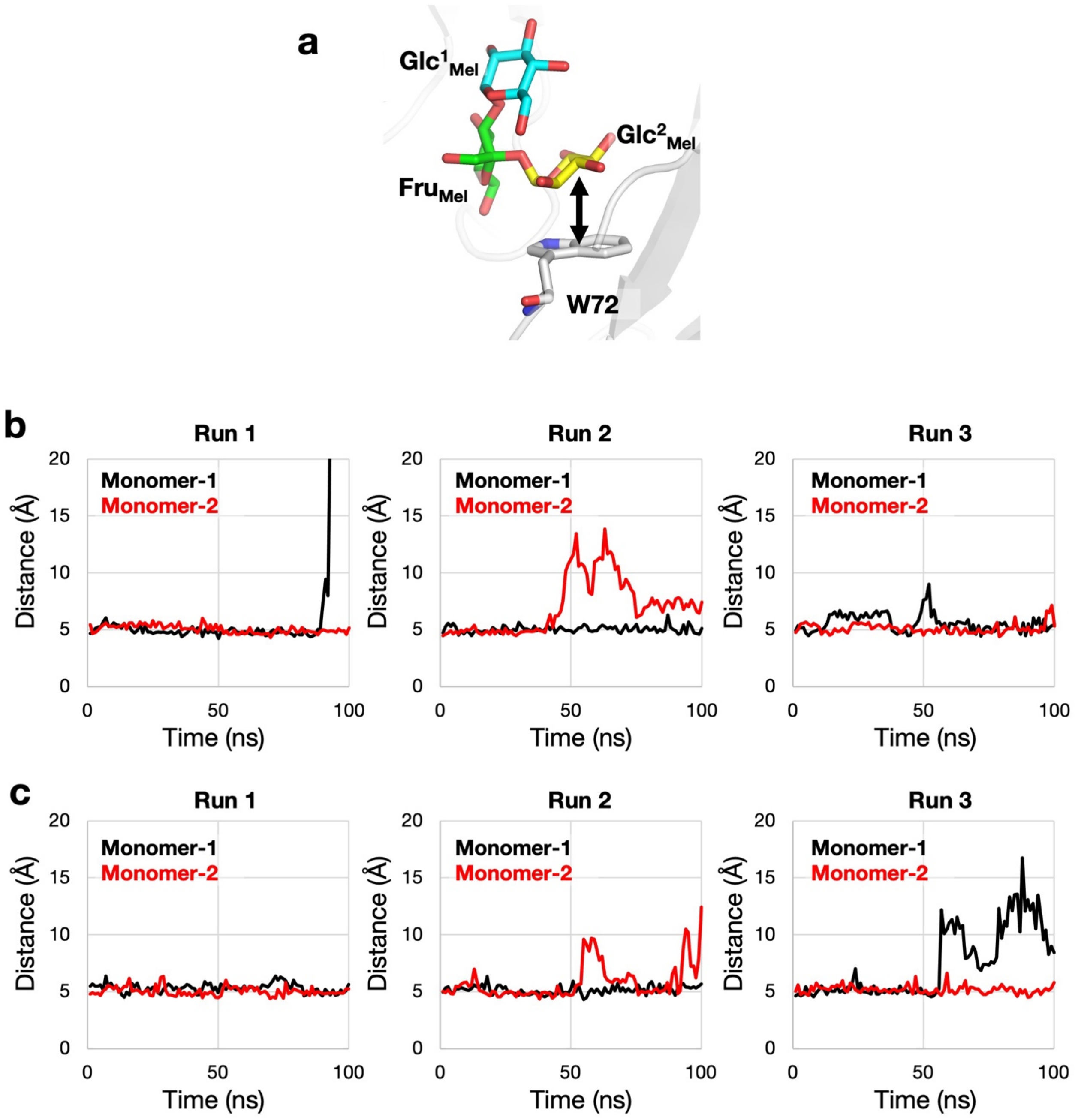


**Figure S8** (a) The close-up views of melezitose in **Mel/E33A-Gal-10$_{\mathbf{cf}}$**. The time course of the two distinct distances between the centroid of W72 and the centroid of the terminal glucose ($Glc^2_{Mel}$) within the dimer (monomer-1 and monomer-2) of (b) dimer and (c) 14-mer during three time 100 ns simulations. The structure of W72 and melezitose are shown as stick models. The structure of proteins is shown as gray-colored ribbon models. The double arrow in (a) shows the distance between the centroid of W72 and the centroid of the terminal glucose ($Glc^2_{Mel}$).

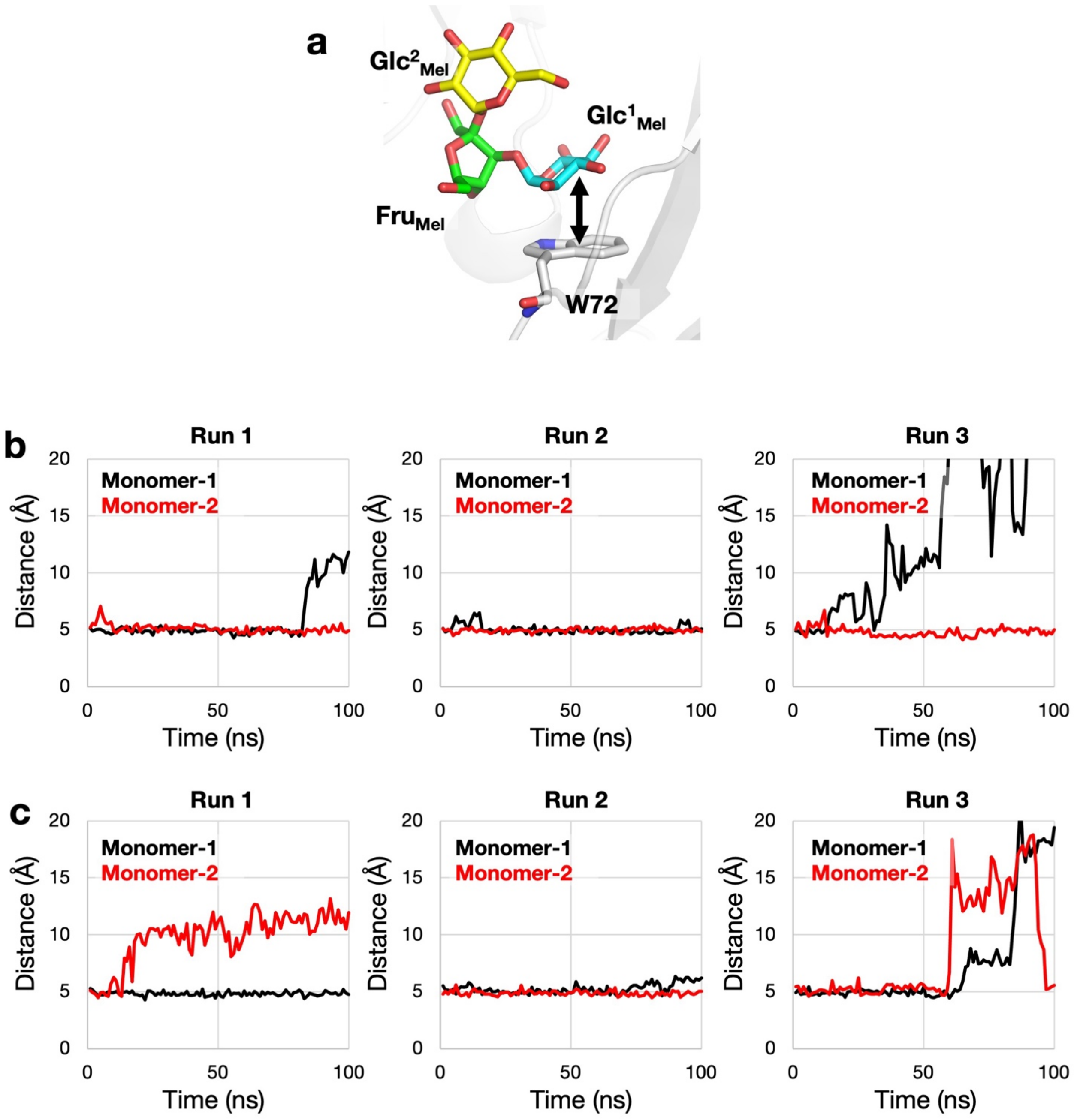


**Figure S9** (a) The close-up views of melezitose in **Mel/Gal-10$_{\mathbf{cf}}$**. The time course of the two distinct distances between the centroid of W72 and the centroid of the terminal glucose (Glc$^{1}_{\mathrm{Mel}}$) within the dimer (monomer-1 and monomer-2) of (b) dimer and (c) 14-mer during three time 100 ns simulations. The structure of W72 and melezitose are shown as stick models. The structure of proteins is shown as gray-colored ribbon models. The double arrow in (a) shows the distance between the centroid of W72 and the centroid of the terminal glucose (Glc$^{1}_{\mathrm{Mel}}$).

**Table S1** Crystallographic data of **Gal-10$_{cf}$** and tri- and disaccharides soaked **Gal-10$_{cf}$s**.

| | **Gal-10$_{cf}$**[a] | **Mel/Gal-10$_{cf}$** | **Raf/Gal-10$_{cf}$** | **Suc/Gal-10$_{cf}$** | **Tre/Gal-10$_{cf}$** | **Mal/Gal-10$_{cf}$** | **Lac/Gal-10$_{cf}$** | **Mel/E33A-Gal-10$_{cf}$** | **Raf/E33A-Gal-10$_{cf}$** |
|---|---|---|---|---|---|---|---|---|---|
| **Soaking condition** | | | | | | | | | |
| Target | – | Melezitose | Raffinose | Sucrose | Trehalose | Maltose | Lactose | Melezitose | Raffinose |
| Concentration | – | 1 M | 0.375 M[a] | 1 M | 1 M | 1M | 0.6 M[a] | 1M | 0.375 M[a] |
| **Data collection** | | | | | | | | | |
| Space group | $P6_522$ | $P6_522$ | $P6_522$ | $P6_522$ | $P6_522$ | $P6_522$ | $P6_522$ | $P6_522$ | $P6_522$ |
| Cell dimensions (Å) | | | | | | | | | |
| *a* | 49.28 | 49.47 | 49.22 | 49.25 | 49.15 | 49.05 | 49.31 | 49.16 | 49.19 |
| *b* | 49.28 | 49.47 | 49.22 | 49.25 | 49.15 | 49.05 | 49.31 | 49.16 | 49.19 |
| *c* | 261.87 | 263.79 | 261.01 | 262.48 | 262.18 | 261.97 | 262.21 | 262.56 | 261.80 |
| Resolution (Å) | 50–1.60 (1.66–1.60) | 50–1.99 (2.11–1.99) | 50–1.81 (1.92–1.81) | 50–1.64 (1.74–1.64) | 50–1.78 (1.89–1.78) | 50–2.31 (2.45–2.31) | 50–1.57 (1.67–1.57) | 50–1.62 (1.72–1.62) | 50–1.57 (1.63–1.57) |
| Observed reflections | 1,414,733 (114,358) | 223,668 (34,406) | 489,823 (776,41) | 1,717,382 (269,836) | 107,511 (17,973) | 45,151 (7,340) | 2,592,922 (338,706) | 366,692 (56,656) | 662,795 (90,384) |
| Unique reflections | 26,219 (2,531) | 14,256 (2,179) | 18,366 (2,853) | 24,512 (3,819) | 18,544 (3,009) | 8,771 (1,391) | 27,804 (4,503) | 25,346 (4,028) | 27,731 (4,553) |
| Redundancy | 54.0 (45.2) | 15.7 (15.8) | 26.7 (27.2) | 70.1 (70.7) | 5.8 (6.0) | 5.2 (5.3) | 93.3 (75.2) | 14.5 (14.1) | 23.9 (2.0) |
| CC(1/2) | 99.9 (77.5) | 98.5 (66.9) | 98.8 (62.4) | 99.1 (63.9) | 99.2 (54.3) | 94.9 (53.2) | 99.5 (77.5) | 98.8 (70.4) | 100 (99.5) |
| I/σ(I) | 18.5 (1.1) | 6.5 (1.7) | 7.8 (1.3) | 23.1 (8.1) | 11.0 (2.7) | 4.1 (1.3) | 32.7 (14.1) | 10.5 (3.0) | 43.5 (12.4) |
| Completeness (%) | 100.0 (100.0) | 99.9 (100.0) | 100.0 (100.0) | 99.9 (99.7) | 96.0 (97.6) | 96.0 (97.8) | 99.6 (99.1) | 99.8 (100.0) | 100.0 (100.0) |
| **Refinement** | | | | | | | | | |
| Resolution (Å) | 33.09–1.60 | 42.29–1.99 | 42.07–1.81 | 42.1–1.64 | 28.12–1.78 | 35.64–2.31 | 42.7–1.57 | 42.02–1.62 | 40.51–1.57 |
| Number of reflections | 26,192 | 14,149 | 17,063 | 24,334 | 18,396 | 8,725 | 27,610 | 25,240 | 27,594 |
| Free *R*-factor (%) | 22.02 | 24.72 | 25.26 | 23.99 | 24.50 | 30.97 | 21.64 | 22.02 | 20.29 |
| R. m. s. deviations | | | | | | | | | |
| Bond lengths (Å) | 0.008 | 0.009 | 0.008 | 0.008 | 0.009 | 0.008 | 0.007 | 0.007 | 0.007 |
| Bond angles (°) | 1.01 | 1.08 | 1.05 | 1.12 | 1.11 | 1.14 | 1.03 | 0.99 | 1.06 |
| Ramachandran plot (%) | | | | | | | | | |
| favoured | 97.83 | 97.10 | 97.08 | 97.81 | 97.12 | 94.89 | 97.83 | 97.83 | 97.84 |
| allowed | 1.45 | 2.17 | 2.19 | 1.46 | 2.16 | 4.38 | 1.45 | 1.45 | 1.44 |
| outlier | 0.72 | 0.72 | 0.73 | 0.73 | 0.72 | 0.73 | 0.72 | 0.72 | 0.72 |
| B-factor (Å$^2$) | | | | | | | | | |
| Protein | 25.33 | 20.99 | 23.89 | 15.57 | 23.01 | 25.79 | 12.83 | 16.16 | 15.75 |
| Saccharide | – | 57.58 | – | 32.97 | 52.40 | 43.58 | 39.34 | 27.04 | 40.40 |
| Monosaccharide 1 | – | 39.66 (Glc$^{1}_{Mel}$) | – | 28.30 (Glc$_{Suc}$) | 40.84 (Glc$^{1}_{Tre}$) | 38.17 (Glc$^{1}_{Mal}$) | 31.47 (Glc$_{Lac}$) | 35.63 (Glc$^{1}_{Mel}$) | 21.65 (Glc$^{1}_{Raf}$) |
| Monosaccharide 2 | – | 63.30 (Fru$_{Mel}$) | – | 38.29 (Fru$_{Suc}$) | 62.92 (Gal$^{2}_{Tre}$) | 48.53 (Glc$^{2}_{Mal}$) | 47.57 (Gal$_{Lac}$) | 30.27 (Fru$_{Mel}$) | 41.03 (Fru$_{Raf}$) |
| Monosaccharide 3 | – | 77.03 (Glc$^{2}_{Mel}$) | – | – | – | – | – | 16.20 (Glc$^{2}_{Mel}$) | 58.52 (Glc$^{2}_{Raf}$) |
| Water | 36.66 | 26.45 | 28.91 | 26.47 | 29.94 | 24.06 | 24.22 | 27.28 | 28.27 |
| **PDB ID** | 9UWK | 9UWU | – | 9UWO | 9UWS | 9UWT | 9UWP | 8JAE | 9UWN |

Values in parentheses are for the highest-resolution shell. [a]Saturated concentration.

### Reference


[1] K. Yamashita, K. Hirata, M. Yamamoto, *Acta Crystallogr. Sect. D: Struct. Biol.* **2018**, *74*, 441-449.

[2] W. Kabsch, *Acta Crystallogr. Sect. D: Biol. Crystallogr.* **2010**, *66*, 133-144.

[3] P. V. Afonine, R. W. Grosse-Kunstleve, N. Echols, J. J. Headd, N. W. Moriarty, M. Mustyakimov, T. C. Terwilliger, A. Urzhumtsev, P. H. Zwart, P. D. Adams, *Acta Crystallogr. Sect. D: Struct. Biol.* **2012**, *68*, 352-367.

[4] S. C. Lovell, I. W. Davis, W. B. Arendall, 3rd, P. I. de Bakker, J. M. Word, M. G. Prisant, J. S. Richardson, D. C. Richardson, *Proteins* **2003**, *50*, 437-450.

[5] T. W. Johnson, R. A. Gallego, A. Brooun, D. Gehlhaar, M. McTigue, *Acs Medicinal Chemistry Letters* **2018**, *9*, 878-883.

[6] W. L. Jorgensen, J. Chandrasekhar, J. D. Madura, R. W. Impey, M. L. Klein, *J. Chem. Phys.* **1983**, *79*, 926-935.

[7] J. A. Maier, C. Martinez, K. Kasavajhala, L. Wickstrom, K. E. Hauser, C. Simmerling, *J. Chem. Theory Comput.* **2015**, *11*, 3696-3713.

[8] K. N. Kirschner, A. B. Yongye, S. M. Tschampel, J. Gonzalez-Outeirino, C. R. Daniels, B. L. Foley, R. J. Woods, *J. Comput. Chem.* **2008**, *29*, 622-655.

[9] R. T. McGibbon, K. A. Beauchamp, M. P. Harrigan, C. Klein, J. M. Swails, C. X. Hernandez, C. R. Schwantes, L. P. Wang, T. J. Lane, V. S. Pande, *Biophys. J.* **2015**, *109*, 1528-1532.

[10] D. Liebschner, P. V. Afonine, N. W. Moriarty, B. K. Poon, O. V. Sobolev, T. C. Terwilliger, P. D. Adams, *Acta Crystallogr. Sect. D: Struct. Biol.* **2017**, *73*, 148-157.

[11] E. N. Baker, R. E. Hubbard, *Prog. Biophys. Mol. Biol.* **1984**, *44*, 97-179.
